# Targeted DNA nicking enables efficient, single-step and counterselection-free editing of bacteriophage genomes

**DOI:** 10.64898/2026.06.25.734514

**Authors:** Frank Englert, Sarshad K. Valappil, Jonas Kubilius, Stephen K. Jones, Vivek K. Mutalik, Chase L. Beisel, Constantinos Patinios

## Abstract

Genetic manipulation of bacteriophages is essential for interrogating phage biology and advancing antimicrobial therapies. However, current genome editing approaches can be inefficient, require multiple steps, or drastically reduce phage titers. Here, we show that targeted DNA nicking enables template-mediated editing of phage genomes in one step without reducing phage titers. Using T7 phage, we show that Cas9-mediated nicking achieved up to 100% recombination across multiple loci, including substitutions and deletions of up to 200 bp and insertions of up to 500 bp, all while preserving phage titers. Editing in T7 was RecA-independent and extended to other phages. Leveraging high titers, we engineered a T7 library of over 440,000 tail-fiber mutants, with isolated mutants restoring infection of two LPS-deficient *Escherichia coli* hosts by shifting recognition to core LPS components. Overall, DNA nicking is a simple and distinct editing strategy that can advance phage genome engineering, genetic interrogation, and antimicrobial development.

## INTRODUCTION

Bacteriophages play a crucial role in the natural world and in biotechnology due to their intrinsic ability to infect, manipulate and kill their bacterial host. They serve as powerful models for understanding fundamental biological processes, such as gene regulation^1,2^, evolution^3,4^, and microbial ecology^5,6^. Moreover, phages are increasingly used in biotechnological applications across human, animal and environmental fields. Their specificity for host bacteria makes them ideal for pathogen control in agriculture^7–9^ and livestock^10,11^, as well as alternative biosensors for the detection of bacteria^12–14^. Phage morphological and genomic plasticity enables them to tolerate modifications without losing functionality, making them highly adaptable for engineering. This flexibility allows technologies like phage display, where foreign peptides or proteins can be presented on the phage surface for applications in drug discovery, diagnostics, and protein engineering^15–17^. To fully harness their potential, it is essential to have efficient access to their genomes and the tools to precisely modify them at single gene resolution.

Gene editing of phages has involved two general approaches distinguished by counterselection. Counterselection-free approaches, such as pORTMAGE and recombitrons, drive editing through heterologous recombinases without inherently selecting against unedited phages^18–21^. These approaches preserve phage titers, although editing efficiencies are often low, requiring multiple rounds of infection and extensive screening to isolate successful mutants. Counterselection-based methods most commonly employ RNA-guided nucleases derived from CRISPR-Cas defense systems programmed to selectively recognize a target DNA or RNA encoded by the phage^22–26^. Target recognition blocks the generation of new phage particles either by cleaving or degrading the DNA target within the phage genome^25–29^ or inducing cell dormancy through widespread collateral RNA cleavage^30^. Successful genome edits prevent CRISPR activation either by disrupting the target sequence or by introducing an expressed anti-CRISPR protein^31^. These edits can be incorporated through homologous recombination mediated by the host or through heterologous expression of recombineering proteins^27,29,32^. These approaches have achieved high editing efficiencies. However, these efficiencies come at the cost of greatly reduced phage titers by eliminating a large number of unedited phages, requiring two-step workflows and reducing the scale of editing that can be performed. The need to block CRISPR activation further restrains the types of edits that can be performed. Therefore, there remains an outstanding need for approaches that offer high editing efficiencies in phages without reducing phage titers or constraining the types of edits.

Here, we report the surprising finding that DNA nicking with Cas9 can mediate efficient and flexible phage editing while maintaining high phage titers. We demonstrated the efficient introduction of deletions, substitutions and insertions to the T7 genome and created a large tail-fiber library with modified receptor binding properties that expanded the phage’s host range to otherwise resistant *E. coli* hosts. Using a genome-wide loss-of-function assay, we further discovered a previously unknown *E. coli* gene (*waaJ*) that confers resistance to some T7 mutant isolates. Finally, we isolated one T7 mutant isolate that exhibited uniform infectivity across all tested LPS *E. coli* mutants. In total, targeted DNA nicking expands the capacity for streamlined and scalable genome editing in phages that can reveal new features of bacterial-phage interactions and facilitate the engineering of phage therapies.

## RESULTS

### Cas9 nickase mediates efficient editing of the T7 phage genome without compromising phage titers

To establish programmable editing of the T7 phage genome, we first tested the previously described CRISPR-Cas9-based genome editing tool involving homologous recombination with a repair template (RT) and selection of mutants through counterselection^33,34^. We used the PAM-flexible (5′-NNG-3′) *Streptococcus canis* Cas9 (ScCas9)^35^, a single-guide RNA (sgRNA) targeting the T7 genome, and a RT encoding the desired edit (5-bp substitution; **Fig. S1**) for the T7 *gp7* gene flanked by 500-bp homology arms. *E. coli* MG1655 and *E. coli* TOP10 were used as hosts, with the latter carrying (among others) a *recA* mutation that prevents efficient host-mediated homologous recombination. Cells were initially transformed with plasmids encoding the ScCas9 nuclease and later with a plasmid carrying the RT and either the targeting (T; sp3) sgRNA for targeting the phage *gp7* gene or a non-targeting (NT) sgRNA serving as control (**Fig. 1A**). Efficient targeting by the sp3 T-sgRNA was verified in phage defense assays, where ScCas9 reduced T7 plaque-forming units (PFU) by ∼1,000-fold compared to the NT-sgRNA control (**Fig. S2**). After plasmid transformation, cells were infected with WT T7 phage and individual plaques were analyzed by Sanger sequencing to assess editing outcomes. The ScCas9^D10A^ nickase (nCas9) and the dead ScCas9^D10A/H848A^ (dCas9) were included as controls, expecting that recombination would be minimal under these conditions.

**Figure 1:**
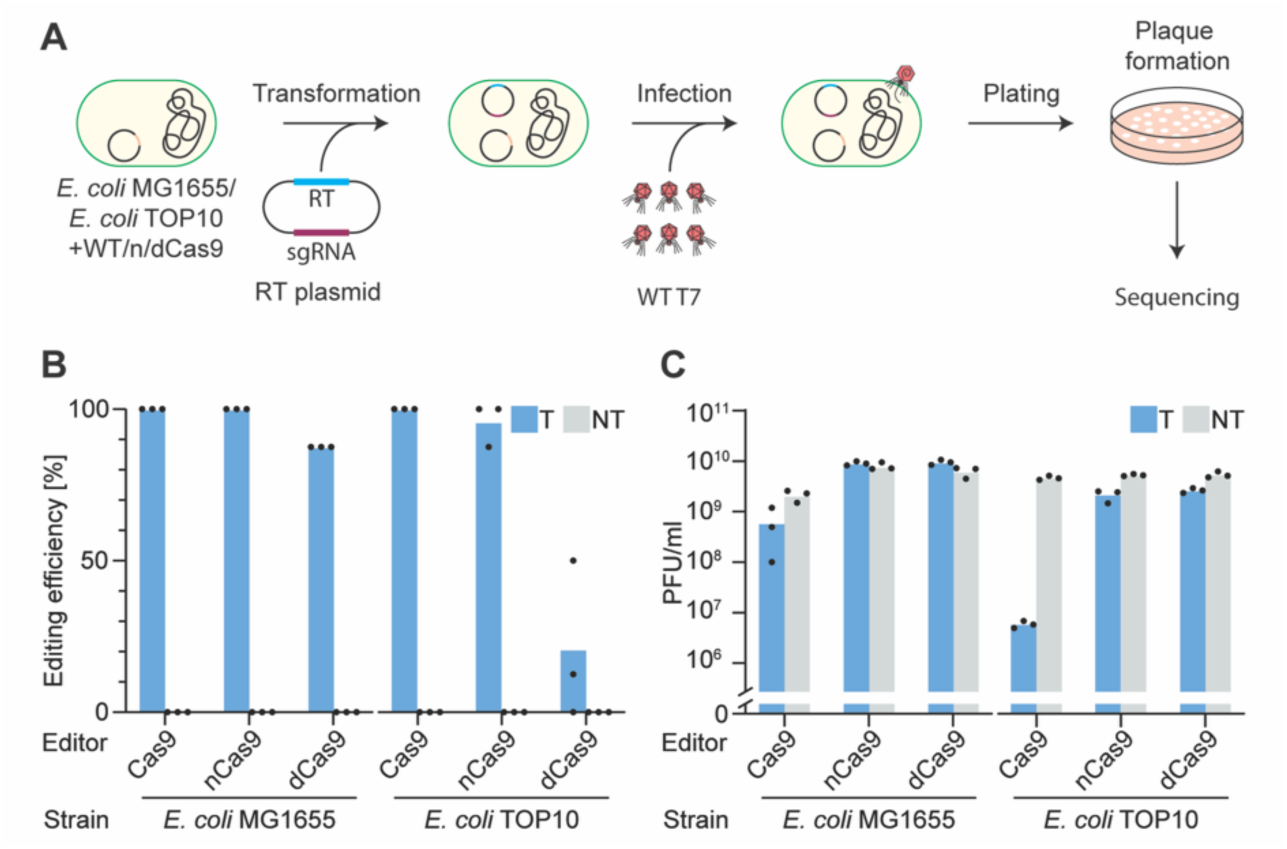
Efficient T7 phage genome editing using Cas9 nickase. **(A)** Workflow to generate phage mutants. *E. coli* cells carrying the nuclease-encoding plasmid are transformed with the RT plasmid containing the sgRNA and the repair template (RT) and then infected with WT T7 phage. The infected cells are plated and the resulting plaques are sequenced through Sanger sequencing. **(B)** Editing efficiency of recovered T7 plaques and **(C)** number of resulting plaque forming units (PFUs) using different Cas9 variants, a targeting (T) or a non-targeting (NT) sgRNA, and TOP10 or MG1655 *E. coli* hosts. Each point in B represents the average editing efficiency from eight plaques screened individually by Sanger sequencing, whereas each point in C represents a biological replicate. Each bar represents the average from three independent biological replicates.

As expected, when using the T-sgRNA, Cas9 yielded 100% editing in both *E. coli* hosts (**Fig. 1B**). Surprisingly, similar editing efficiency was also observed with nCas9 in both hosts. Furthermore, dCas9 resulted in 87% editing efficiency in *E. coli* MG1655 and ∼20% in *E. coli* TOP10. Plaque formation was maintained between 1.50 × 10^9^ and 1.06 × 10^10^ under targeting and non-targeting conditions for both nCas9 and dCas9 in either host (**Fig. 1C**). In contrast, Cas9 combined with a T-sgRNA reduced plaque formation by ∼3.5-fold in *E. coli* MG1655 and by ∼800-fold in *E. coli* TOP10 compared to the NT-sgRNA control. Control conditions using a NT-sgRNA combined with any of the tested nucleases showed no detectable edits across both host strains (**Fig. 1C**). Combined, these results show that DNA nicking with Cas9 ickase mediates high editing efficiency in both *E. coli* hosts without a considerable drop in phage titers.

### nCas9 enables flexible genome editing in phages

To gauge the potential of nCas9 for phage genome editing, we evaluated the impact of homology arm length on editing outcomes. By following the same workflow as in **figure 1A** and by using Oxford Nanopore sequencing as the last step for editing efficiency quantification, we systematically shortened the size of the RT. When 500 and 200 bp long homology arms were used, >90% editing efficiency was observed (**Fig. 2A**). Further shortening of the length of the homology arms led to a sharp drop in editing efficiency that became negligible below 50 bp. Phage titers varied across the tested homology arm lengths but stayed consistently high (>10^9^) throughout the tested conditions (**Fig. 2A bottom**). Using 500-bp homology arms, we created substitutions up to 200 bp with editing rates of ∼70%, deletions up to 200 bp with almost 100% editing efficiency, and insertions up to 100 bp with over 80% editing efficiencies and 500 bp with ∼50% editing efficiency (**Fig. 2A**). Phage titers showed some variance depending on the type and size of edit but stayed consistently above 10^8^ PFU/ml (**Fig. 2B bottom**).

**Figure 2:**
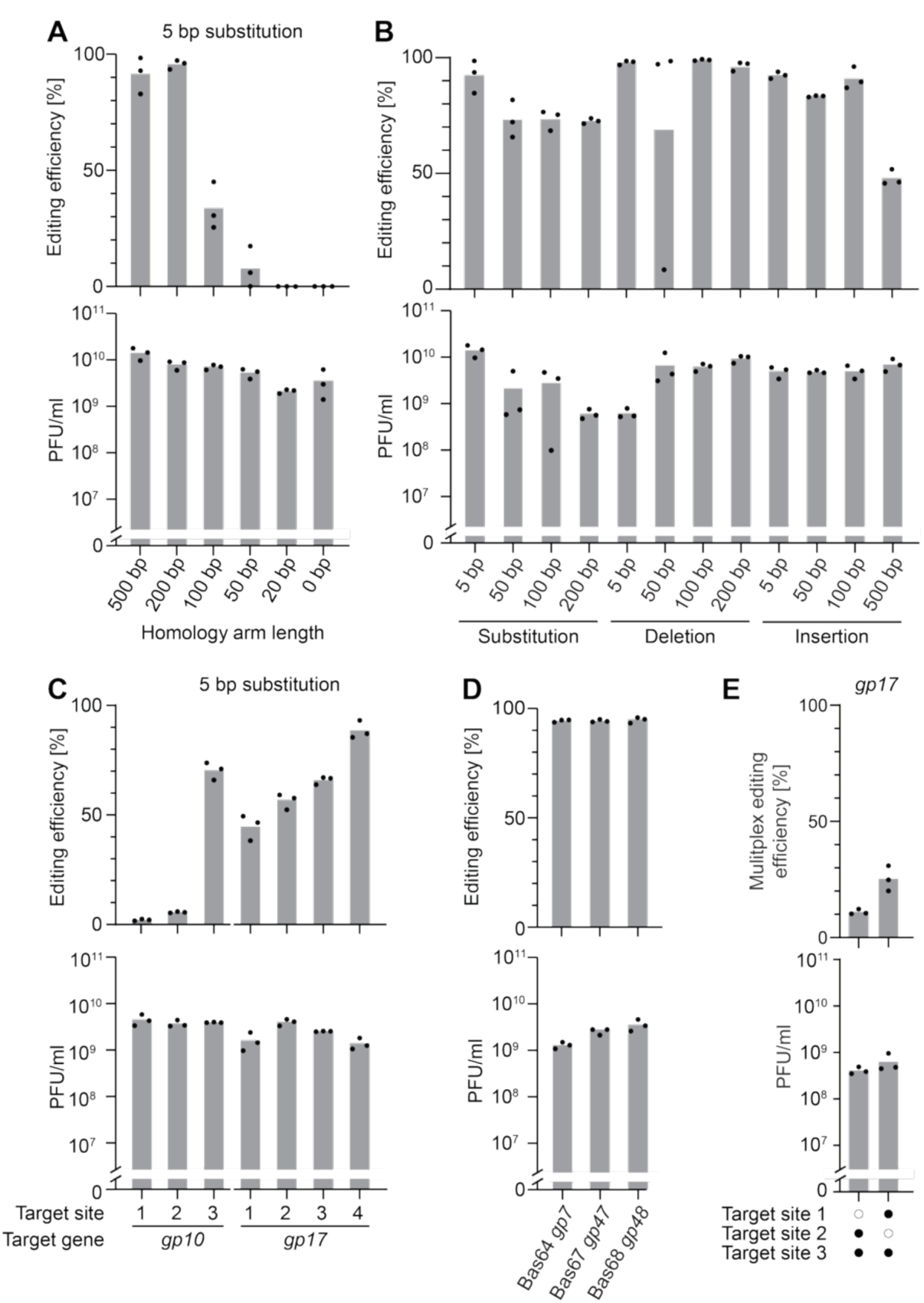
Flexible genome editing in *E. coli* phages using nCas9. **(A)** Editing efficiency (top) and phage titer (bottom) for 5 bp substitutions in the T7 phage *gp7* gene, using repair templates with homology arm lengths ranging from 0 bp to 500 bp. **(B)** Editing efficiency (top) and phage titer (bottom) for 5-500 bp substitutions, deletions and insertions in the phage T7 *gp7* gene. **(C)** Editing efficiency (top) and phage titer (bottom) for 5 bp substitutions at different sites within the T7 phage *gp10* and *gp17* genes. **(D)** Editing efficiency (top) and phage titer (bottom) for 10 bp substitutions in phages Bas64, Bas67, and Bas68. **(E)** Editing efficiency (top) and phage titer (bottom) for two simultaneous 5-bp substitutions in the T7 phage *gp17* gene during a single round of editing. Each bar represents the average from three independent biological replicates. Each dot represents a biological replicate. Editing was defined through Oxford Nanopore sequencing.

To explore genome editing with nCas9 beyond the *gp7* gene in T7, we targeted *gp10* and *gp17*, encoding for the capsid protein and the tail fiber protein, respectively. For one out of three target sites in *gp10* (target site 3), we introduced 5 bp substitutions in ∼70% of screened phages, while the other two targets resulted in <10% editing efficiency (**Fig. 2C**). Targeting four different sites within *gp17*, we achieved editing between ∼40% and ∼90%. Across all seven tested target sites, phage counts were consistently >10^9^ PFU/ml (**Fig. 2C bottom**).

We further expanded the applicability of our phage-editing tool by targeting additional *E. coli* phages. Because editing remained efficient in *E. coli* TOP10 despite the absence of RecA, we speculated that phage-encoded factors, such as the essential Gp2.5 protein^36–38^, promote homologous recombination following ssDNA nick formation. We therefore screened the BASEL collection for phages encoding homologous proteins to T7 Gp2.5 and selected Bas64, Bas67, and Bas68 for editing (**Fig. S3**). The corresponding proteins shared 98.7%, 62.7%, and 61.9% sequence identity with T7 Gp2.5, respectively. We introduced 8-10 bp substitutions in the genomes of Bas64, Bas67 and Bas68, with editing frequencies exceeding 90% and phage titers close to or above 10^9^ PFU/ml (**Figs. 2D and S4**).

To simplify and accelerate editing at multiple genomic loci, a previous study introduced multiple edits into lambda (Δ*cI*) phage using a mixture of bacterial strains, each carrying a distinct editing recombitron^21^. Building on this mixed-host strategy, we assessed its compatibility with nCas9-mediated phage genome editing by combining *E. coli* strains carrying the same nCas9 plasmid but different RT plasmids, followed by simultaneous infection with WT T7 phage (**Fig. S5**). For this, we chose the previously analysed targets within *gp17* due to their close genomic location, allowing us to easily identify if phage genomes contained individual or cumulative edits through Oxford nanopore sequencing (**Fig. S6**). Combining *E. coli* hosts that carried RT plasmids targeting target sites 2 and 3 or 1 and 3, we were able to identify multiplex genome editing in ∼10% and ∼20% of phage genomes, respectively, with phage titers maintained at >10^8^ PFU/ml **(Fig. 2E)**. Analysis of individual edits showed that editing frequencies varied between ∼10% and ∼60%, depending on the target site and the combination of targets **(Fig. S7)**.

### Large mutant library generation at the T7 *gp7* and *gp17* genes

Given the high editing efficiency and the consistent high phage titers, we asked whether we could generate large mutant libraries at designated genomic locations on the T7 genome. As a proof of principle, we followed the workflow indicated in **figure 3A**, and targeted *gp7*, a non-essential gene that we have edited successfully (**Fig. 2C**). We chose to substitute 12 nucleotides at the protospacer 3 site with every possible nucleotide combination, resulting in a theoretical library of 4^12^ (1.7 × 10^7^) variants (**Fig. 3B**). We recovered 1.5 ×10^9^ PFUs/ml, with ∼75% of the recovered phages edited at the target site (**Fig. 3C and Fig. S8**). Through NGS, we were also able to determine that our screened replicates showed ∼85,000 and ∼97,000 unique sequences each, resulting in ∼180,000 (1.8 × 10⁵) unique sequences when combined, representing ∼1% of the theoretical mutant library size (**Fig. 3D**).

**Figure 3:**
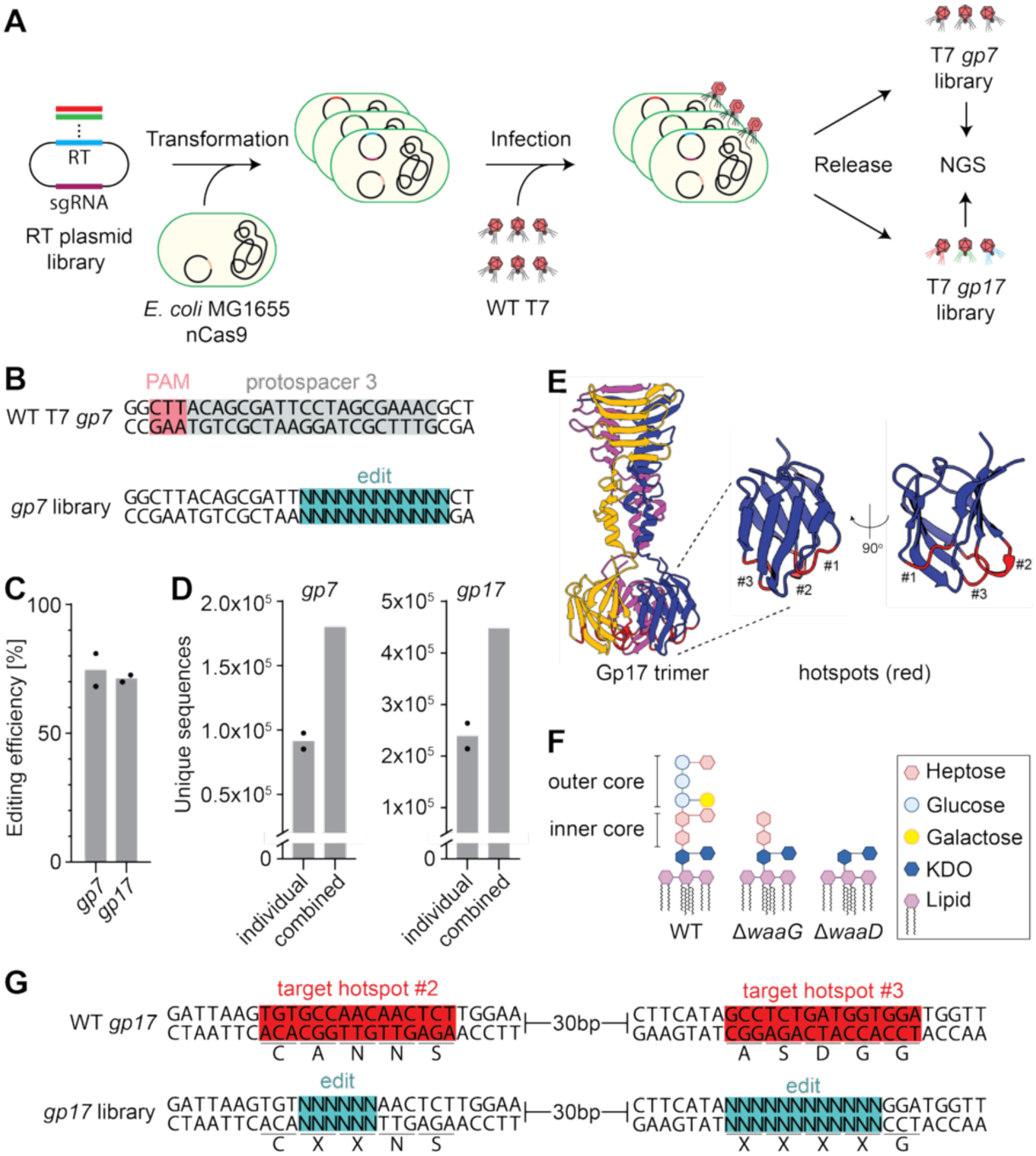
nCas9 enables the engineering of large T7 mutant libraries. **(A)** Workflow for generating T7 phage *gp7* and *gp17* mutant libraries. The repair template (RT) plasmid library was transformed into *E. coli* MG1655 carrying the nCas9 plasmid. The host cells were then infected with wild-type (WT) T7 phages, and the released phages were analyzed by next-generation sequencing (NGS). **(B)** Overview of the T7 *gp7* gene targeted protospacer sequence highlighted in grey, with the ScCas9 PAM highlighted in pink and the introduced 12 × N library region highlighted in teal. **(C)** Editing efficiency as determined by NGS for the T7 *gp7* and *gp17* genes. **(D)** Unique sequence counts for the individual and combined library duplicates of *gp7* (left) and *gp17* (right) genes. **(E)** Structure of the T7 tail fiber (Gp17 trimer) with a close-up of the outer loops at the tip showing the three hotspots in red. **(F)** Schematic overview of the LPS composition of WT, Δ*waaD* and Δ*waaG E. coli* strains. **(G)** Overview of the targeted hotspot sequences of the T7 *gp17* gene highlighted in red and the introduced 6 × N and 12 × N libraries highlighted in teal. The encoded amino acids are denoted below.

Encouraged by our ability to generate hundreds of thousands of unique mutants, we used nCas9 to target and mutate the T7 *gp17* gene which encodes for the tail fiber protein that mediates adsorption to the host by binding the primary receptor^39^ (**Fig. 3E**). The exposed outer loops of *gp17* are responsible for the initial binding to the host lipopolysaccharide (LPS) and are essential for the adsorption of T7 to their host^40^.

Alterations to the host LPS, render the adsorption impossible or inefficient^41,42^. Examples include the deletion of the *waaD* or *waaG* genes that result in shortening the *E. coli* LPS and confers resistance to T7 infection^43,44^ (**Fig. 3F**). *waaD* encodes an ADP-L-glycero-D-manno-heptose-6-epimerase, which attaches ADP-heptose residues to the 3-deoxy-D-manno-oct-2-ulosonic acid (Kdo) of the inner core. *waaG* encodes for a glucosyltransferase, which catalyzes the addition of the first outer-core glucose to the inner core. In Δ*waaD* and Δ*waaG* mutant strains, the LPS is therefore truncated at the inner core at the Kdo and Heptose residues, respectively.

Prior work showed that by modifying the Gp17 protein at specific “hotspot” regions (the exposed outer loops), mutant T7 isolates were able to infect the Δ*waaD* and Δ*waaG E. coli* strains, overcoming host incompatibility based on tail fiber-LPS interactions^39,44^. We randomized two of the previously identified hotspots (#2 and #3), mutating six nucleotides (i.e., two amino acid residues) of hotspot #2 and twelve nucleotides (i.e., four amino acid residues) of hotspot #3 (**Fig. 3G**). In total, randomizing these 18 positions had the potential of creating 4^18^ (or ∼10^11^) possible genetic variants. Infection of the *E. coli* host with WT T7 phage and subsequent plating yielded ∼5 × 10^9^ PFU/ml (**Fig. S8**). The recovered pooled phage library was sequenced through NGS, revealing ∼70% editing (**Fig. 3C**) and ∼260,000 and ∼210,000 unique sequences present in each of our two replicates. Very few of these unique sequences were shared between the two replicates, forming a library of ∼440,000 unique sequences (**Fig. 3D**).

### Tail fiber library allows enrichment of T7 mutants on otherwise resistant *E. coli* hosts

To test whether the introduced library mutations at the Gp17 can expand the host range, we infected *E. coli* MG1655 Δ*waaD* and Δ*waaG* hosts with the *gp17* mutant library, collected the progeny phages (referred to as waaDi and waaGi; i standing for isolates) and sequenced the *gp17* hotspot #2 and #3 regions using NGS (**Fig. 4A**). ∼95% of the waaDi and ∼60% of the waaGi were edited at least at one of the two hotspots (**Fig. 4B**). Both replicates of waaDi and waaGi pools had ∼30,000 and ∼29,000 unique sequences, respectively, and ∼57,000 and ∼55,000 unique sequences when combined (**Fig. 4C**). In waaDi replicate 1, the WT *gp17* sequence ranked 12th in abundance and accounted for ∼2% of total reads, whereas in replicate 2 it ranked 3rd and represented ∼9% of total reads (Fig. S9). By contrast, the WT *gp17* sequence was the most abundant variant in both waaGi replicates, comprising ∼42% and ∼30% of total reads in replicates 1 and 2, respectively (**Fig. S9**).

**Figure 4:**
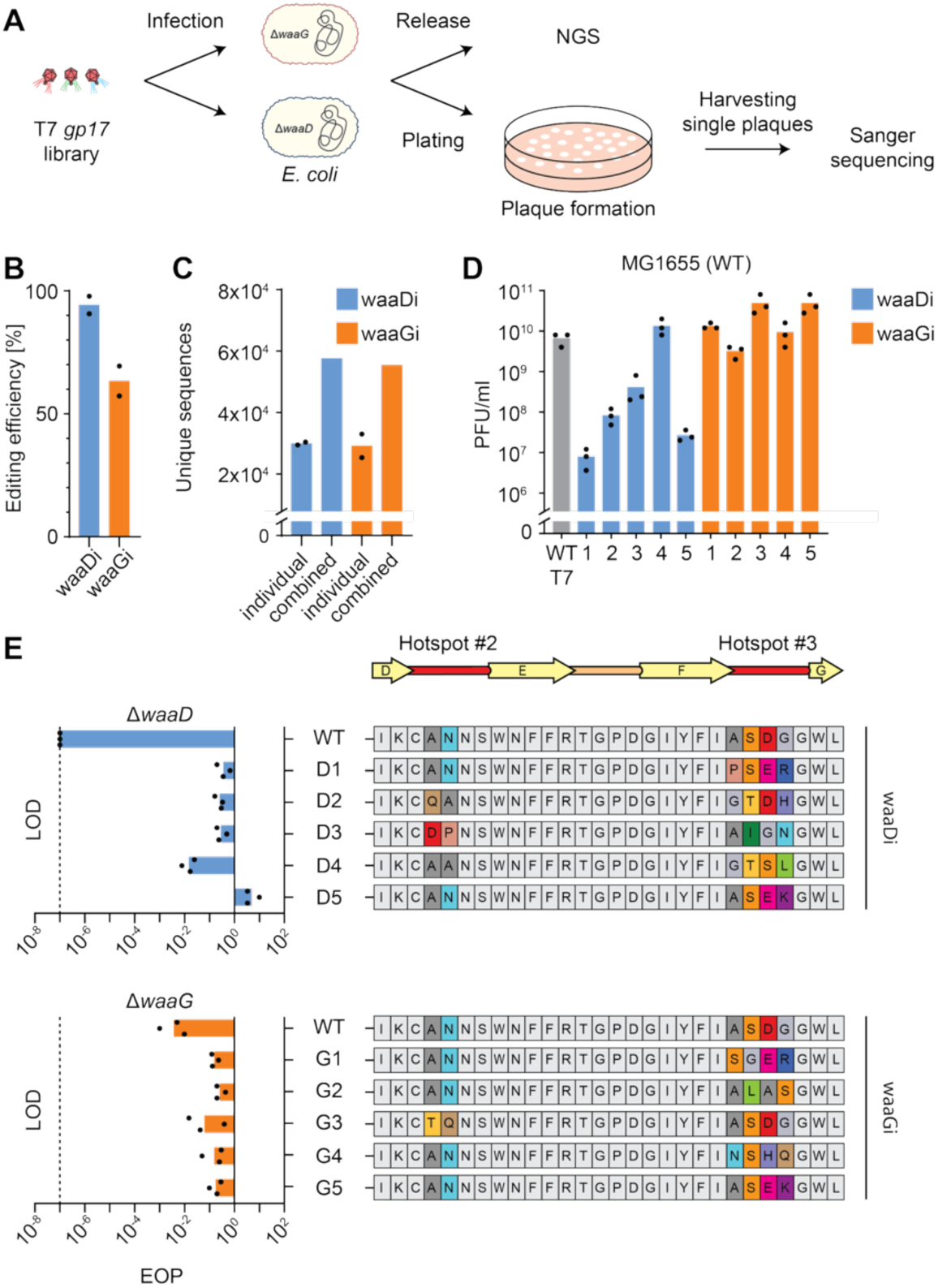
Engineered tail fiber libraries allow enrichment of T7 mutants on otherwise resistant *E. coli* hosts. **(A)** Workflow for enrichment of the T7 *gp17* mutant library in mutant *E. coli* hosts. The isolated T7 *gp17* library was used to infect the Δ*waaD* and Δ*waaG E. coli* LPS mutants. After plating, individual plaques were picked and screened through Sanger sequencing. Afterwards, all remaining plaques were harvested and screened through NGS sequencing. **(B)** Editing efficiency as determined by NGS for the mutant phages harvested from the corresponding Δ*waaD* and Δ*waaG E. coli* mutants. The “i” next to waaD or waaG refers to phage isolates harvested from Δ*waaD* or Δ*waaG E. coli* mutants. **(C)** Unique sequence counts for the individual and combined waaDi and waaGi mutant T7 library duplicates. **(D)** Phage titers of WT T7 phage and five waaDi and waaGi, determined on WT *E. coli* MG1655. **(E)** Efficiency of plating (EOP) of WT phage T7 and five waaDi (top) or five waaGi (bottom), determined on *E. coli* Δ*waaD* (top) and Δ*waaG* (bottom). The relevant amino acid sequence of *gp17* hotspot #2 and #3 is shown on the right, and the residues targeted by the libraries are colored. The Gp17 outer loops (red) connecting the parallel β-sheets (D-G, yellow) are shown above. The bars of B and C represent the average of two independent biological replicates, while the bars of D and E represent the average of three independent biological replicates.

Analysis of the recovered variants revealed distinct residue preferences at the two engineered *gp17* hotspots (**Fig. S10**). At hotspot #2, waaDi phages retained the WT alanine at position 1 or substituted it with asparagine, while the WT asparagine at position 2 was retained or replaced by proline or methionine. WaaGi phages showed greater variability but predominantly retained the WT alanine-asparagine sequence. At hotspot #3, both waaDi and waaGi phages largely conserved the WT serine and aspartate at positions 2 and 3, whereas positions 1 and 4 were more variable, with methionine, asparagine, threonine, or proline at position 1 and arginine or serine at position 4. To determine whether per-position preferences are sufficient to predict variant efficacy, we modeled efficacy with a non-linear regression allowing epistatic interactions between positions. This model predicted the efficacy of unseen variants far better than considering residues independently (i.e., sequence logo; Spearman r = 0.16/0.16 vs 0.60/0.54 for waaDi/waaGi). This indicates that variant efficacy is governed by residue combinations rather than conserved positions alone and can be predicted from sequence (**Fig. S11**).

We next asked whether individual or combined mutations in the T7 Gp17 tail fiber altered overall infectivity. To investigate this, we isolated five individual plaques from the waaDi population (D1–D5) and five from the waaGi population (G1–G5) and performed plaque assays using WT, Δ*waaD*, and Δ*waaG E. coli* MG1655 hosts, checking for phage titers and EOP (**Fig. 4D, E and Fig. S12**). Most of the waaDi had ∼10 to 1000-fold reduced PFUs/ml when tested against the WT MG1655 host compared to WT T7 phage, while the waaGi maintained similar infectivity as the WT T7 phage (**Fig. 4D**). The efficiency of plating (EOP) of the waaDi phages was restored against the Δ*waaD* host, with over 150,000-fold improvement compared to WT T7 phage (**Fig. 4E and Fig. S12**). Notably, the waaDi phages maintained an EOP between 1.66 × 10^-2^ and 5.56 × 10⁰, indicating that Gp17 amino acid changes likely contribute to the recognition of an LPS component shared between the WT and Δ*waaD E. coli* strains (**Fig. 4E**). The EOP of the waaGi phages improved by 28- to 52-fold relative to WT T7 (**Fig. 4E and Fig. S12**). Collectively, our results show that nCas9 can create large mutant libraries in a single step at two different hotspots of the *gp17* gene, restoring infectivity on otherwise resistant hosts.

### RB-TnSeq identifies *waaJ* as a key host gene required for efficient mutant phage infection

Enrichment of T7 mutants on either the Δ*waaD* or Δ*waaG E. coli* MG1655 hosts resulted in phages capable of infecting their respective mutant strain. To identify the genetic factors underlying infection of these mutant strains, we used the RB-TnSeq assay that rapidly probes phage-host interaction mechanisms, revealing host factors that contribute to successful phage infection^45^.

RB-TnSeq screening of the ten T7 *gp17* tail fiber mutant isolates (D1-5 and G1-5) revealed a fitness landscape markedly distinct from that of the WT T7 phage (**Fig. 5A**). WT T7 exhibited strong signals of high fitness across several genes involved in LPS biosynthesis, including *waaC*, *waaD*, *waaE*, *waaJ*, and *lpcA*. This reflects its established dependence on a fully assembled LPS core as genes that are critical for Kdo and inner-core heptose biosynthesis scored high fitness scores. Six of the ten *gp17* mutant isolates displayed a narrower fitness profile compared to the WT T7 phage. Isolates D2, D4, G1-3, and G5 were affected mainly by the *waaJ* gene encoding for UDP-glucose:(glucosyl)LPS alpha-1,2-glucosyltransferase, represented by a high fitness score (>7). G1 and G3 were also affected by *lpcA* having a fitness score of 4.8 and 6.4, respectively. Isolates D1, D3, D5, and G4 did not conform to this pattern. They yielded inconclusive fitness signals in the RB-TnSeq screen, suggesting that these mutants may be able to engage inner-core LPS structures or simultaneously utilize multiple sugar moieties during host adsorption, or rely on features created by essential genes.

**Figure 5:**
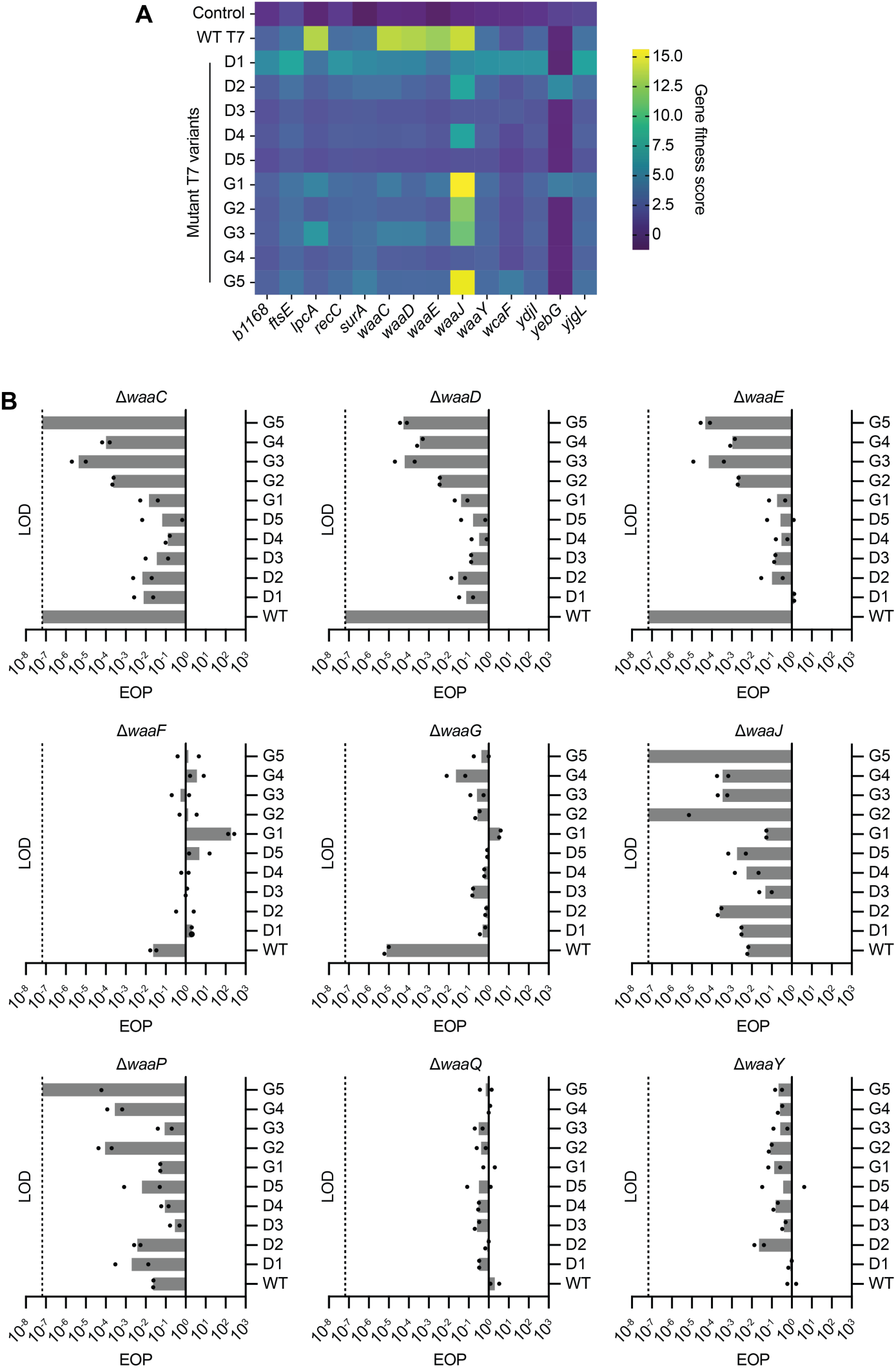
Host range determinants, defined by RB-TnSeq and EOP assays. (**A**) Heatmap of *E. coli* K-12 RB-TnSeq data comparing the T7 phage mutants selected on Δ*waaD* (D1-D5) and on Δ*waaG* (G1-5) *E. coli* hosts, to the WT T7 phage at a multiplicity of infection (MOI) of 1. The top 14 genes showing high-confidence effects, with a gene fitness score greater than 6 in at least one phage assay, are shown. (**B**) EOP of the WT T7 phage and the five isolates selected on Δ*waaD* (D1-D5) and on Δ*waaG* (G1-5) *E. coli* hosts, against nine defined LPS mutant hosts (Δ*waaC*, Δ*waaD*, Δ*waaE*, Δ*waaF*, Δ*waaG*, Δ*waaJ*, Δ*waaP*, Δ*waaQ*, and Δ*waaY*). All EOP assays were performed in duplicate. LOD: Limit of detection.

To functionally validate the RB-TnSeq fitness predictions, EOP assays were performed with phage isolates D1–5 and G1–5, and the WT T7 phage on nine LPS mutant hosts available in the KEIO collection (Δ*waaC*, Δ*waaD*, Δ*waaE*, Δ*waaF*, Δ*waaG*, Δ*waaJ*, Δ*waaP*, Δ*waaQ*, and Δ*waaY*), using WT *E. coli* BW25113 as the reference control (**Fig. 5B and Fig. S13-15**). As expected, WT T7 phage was unable to infect the Δ*waaC*, Δ*waaD*, and Δ*waaE* mutant *E. coli* hosts, confirming our obtained data through RB-TnSeq and the dependence of WT T7 phage on an intact LPS. In contrast to our RB-TnSeq data, the Δ*waaC*, Δ*waaD*, and Δ*waaE* mutant hosts were also effective against the G2-5 isolates with an EOP <1.00 × 10^-3^. However, isolates D1-D5 and G1, experienced moderate inhibition by these mutant hosts, having an EOP of 1.3 × 10^-2^ or higher.

The G5 isolate was unable to infect the Δ*waaJ* host whilst D2, G3 and G4 isolates had an EOP equa to ∼10^-4^. The rest of the isolates experienced a moderate effect. Surprisingly, isolate G1 and the WT T7 phage showed 5.33 × 10^-2^ and 6.33 × 10^-3^ EOP, respectively, in contrast to the high fitness gene score observed during the RB-TnSeq assays. Noticeable inhibition was also observed by the Δ*waaP* host for isolates G2, G4 and G5 but not for other isolates or the WT T7 phage. This gene was also not identified by the RB-TnSeq assay. The hosts Δ*waaF*, Δ*waaQ*, and Δ*waaY* did not have any inhibitory effect for any of the isolates or the WT T7 phage.

Whilst most of the G isolates experienced a narrower host range (except for G1), the D isolates showed good infectivity across all nine LPS mutant hosts. A notable isolate, D3, showed the best overall host range infectivity with the lowest EOP reaching 6.17 × 10^-2^ when infecting the Δ*waaJ* host. This is not a surprise as D3 along with the other waaDi phages have been selected on a minimal LPS mutant, Δ*waaD*, and are likely binding the Kdo or the lipid surface itself. The broad infectivity profiles of these isolates help explain the inconclusive results from the RB-TnSeq signals. A phage that is unaffected by the outer or inner core LPS modifications is unlikely to produce a defined fitness peak at any single locus within a transposon library screen.

## Discussion

Our study establishes targeted DNA nicking as an efficient and single-step genome editing approach for some bacteriophages, enabling precise substitutions, deletions, and insertions without compromising phage titers. By demonstrating editing across multiple loci, different phages, and the capacity to generate large mutant libraries, we highlight the versatility and robustness of this tool.

Compared to existing phage-editing approaches, nCas9 offers several advantages. By introducing single-strand DNA nicks rather than the double-strand breaks generated by Cas9 or Cas12a, nCas9 enables efficient editing without counterselection while preserving high phage titers. This feature is particularly advantageous for large-scale library applications, in which preserving phage titers is essential for maintaining library diversity and adequate variant representation. Moreover, Cas13a-based systems rely on homologous recombination followed by counterselection through target-RNA recognition and collateral RNA cleavage, restricting their use to transcribed genomic regions^30,31^. An alternative could be the integration of an anti-CRISPR (Acr) at the specific genomic location, to counteract the activity of Cas13a. However, this will always necessitate the integration of an Acr, restricting the wide applicability and scarless use of this approach. By contrast, nCas9 acts directly on the phage genome by introducing a site-specific DNA nick and can therefore target both transcribed sequences and non-transcribed regulatory regions, including promoters. Other dsDNA break-free methods, such as recombitrons, are becoming increasingly used, although they currently suffer from low editing efficiency, they require the expression of additional recombineering proteins, they require multiple rounds of infection to enrich edited phages, and large scale mutagenesis has not been demonstrated yet^20,21^. Still, nCas9 has several limitations, including the limited ability of DNA-targeting nucleases to access chemically modified phage genomes^46,47^, uncertainty about how well this mode of editing can be achieved across the diversity of phages, and variable editing efficiencies across genes, loci, and phages. Future work could therefore focus on expanding and improving nicking-based phage editing across phages.

Although our findings are consistent with template-directed repair at the nicked site, the underlying molecular mechanism remains unresolved. nCas9-mediated editing remained highly efficient in both *E. coli* TOP10 and MG1655 without reducing phage titers. By contrast, Cas9 caused an approximately 100-fold reduction in phage titers in TOP10 than in MG1655, potentially reflecting the RecA deficiency of TOP10 and the importance of RecA in homologous recombination and double-strand-break repair. Nevertheless, comparable editing efficiencies were obtained with Cas9 and nCas9 in both strains, suggesting that phage-encoded recombination pathways may also contribute to donor incorporation. One possible contributor is the T7 single-stranded DNA-binding protein Gp2.5, which promotes annealing of complementary DNA strands and participates in RecA-independent homologous recombination and double-strand-break repair^36,37,48–51^. Although gp2.5 has not been directly implicated in the repair of isolated DNA nicks, it could facilitate donor incorporation by annealing single-stranded intermediates generated during nick processing or replication. Direct genetic testing is challenging because *gp2.5* is essential for T7 DNA synthesis, and its inactivation prevents productive phage replication^38^. Moreover, because T7 DNA polymerase can bypass DNA nicks^52^, it remains unclear how nicking is converted into such an efficient recombination substrate. Efficient editing with preserved phage titers was also observed for Bas64, Bas67 and Bas68, indicating that genome editing through DNA nicks is not limited to T7. However, whether these phages use analogous repair pathways, and whether nick-mediated editing is broadly applicable across diverse phages, remains unknown and will likely depend on the recombination and replication machinery encoded by each phage.

As a proof of principle for large-scale library generation, we used nCas9 to diversify the T7 tail-fiber gene *gp17* and selected variants capable of infecting otherwise resistant *E. coli* Δ*waaG* and Δ*waaD* hosts. Combined RB-TnSeq and EOP analyses revealed distinct changes in receptor usage. For most recovered T7 variants, *waaJ* was the dominant determinant of susceptibility, suggesting that *gp17* diversification redirected recognition toward the outer-core glucose added by WaaJ. This differs from wild-type T7, which relies on a more complete LPS core involving genes such as *waaC*, *waaD* and *waaE*, and demonstrates that large-scale tail-fiber mutagenesis can rapidly overcome resistance caused by alterations in LPS biosynthesis. Whereas several Δ*waaG* isolates showed narrower dependencies on specific LPS structures, Δ*waaD* isolates retained infectivity across more extensively truncated LPS backgrounds, indicating reduced reliance on the extended core. The D3 isolate displayed the broadest phenotype, maintaining infectivity across all tested LPS *E. coli* mutants, with only a moderate reduction on Δ*waaJ*. This suggests that D3 may recognize a conserved and broadly accessible surface feature, such as Kdo residues or structures near lipid A. However, its receptor cannot be identified without direct adsorption, competition, or structural studies. In total, high diversity *gp17* mutagenesis generated both variants with redirected receptor specificity and variants such as D3 with reduced receptor stringency. The latter variants are particularly relevant to phage therapy because they may remain infectious as bacteria may progressively modify or lose their LPS structures^41,42,53^.

In summary, this work demonstrates how DNA nicks can facilitate efficient editing of phage genomes without compromising phage titers, allowing large phage mutagenesis. In turn, this approach can be leveraged both for mechanistic insights into phage biology and for developing tailored phages with potential applications in biotechnology and phage therapy, where limited host range and emergence of resistance remain major challenges.

## Methods

### Bacterial strains and growth conditions

*E. coli* strains used in this study (**supplementary table S1**) were cultivated at 37°C in LB medium (10 g/L Tryptone, 5 g/L NaCl and 5 g/L yeast extract) or grown on LB agar (Carl Roth, Cat. # X965.3) plates, supplemented with 34 µg/ml chloramphenicol (Cm; Carl Roth, Cat. # 3886.2), 100 µg/ml carbenicillin (Carb; Carl Roth, Cat. # 6344.3) or 50 µg/ml kanamycin (Kan; Carl Roth, Cat. # T832.3), when appropriate. To induce expression from the arabinose inducible promoter, 0.5 g/L L-arabinose (Sigma-Aldrich, A3256-25G) was added in the growth medium.

### Phage propagation

Depending on the phage (**supplementary table S1**), *E. coli* MG1655 or BW25113 host strains were grown in 3 ml LB medium at 37°C, shaking orbitally at 220 rpm until they reached an OD_600_ equal to 0.3. The bacterial culture was infected with either 10 µL of phage lysate or with a small amount of cryopreserved phages. The infected cultures were incubated 10 min at 37°C and afterwards moved to a shaker continuing the incubation while shaking at 220 rpm. After 3 h or after the culture had cleared, 1.8 ml of the lysate was aliquoted and mixed with chloroform (ITW Reagents, Cat. # 131252.1612) to reach a 0.5% (v/v) final chloroform concentration. After 1 min centrifugation at 11,000 xg, the upper phase was filtered through a 0.2 µm filter (Sarstedt, Cat. # 83.1826.001) and stored at 4°C until further use.

### Plaque assay for phage quantification

For plaque assays to quantify phage titer, the double agar overlay method was used^54^. Individual colonies of host bacteria were cultured in 3 ml of LB medium and incubated at 37°C, shaking orbitally at 220 rpm, for 16 h. The cultures were back diluted in 5 ml LB medium to OD_600_ equal to 0.05 and incubated at 37°C, shaking orbitally at 220 rpm until they reached an OD_600_ equal to 0.5. 2 ml of the bacterial culture was then pelleted at 5000 xg for 3 min and resuspended in 200 µL LB + Cm + Carb medium. 100 µL of this bacterial suspension was mixed with 3 ml pre-warmed LB, 0.7% (w/v) agar (Th.Geyer, Cat. # 214520), vortexed and poured on top of LB agar plates. The tested phages were serially diluted in SM-buffer (5.8 g/L NaCl, 2 g/L MgSO_4_, 50 mM Tris-HCl pH 7.4) and 3 µL droplets were spotted on the solidified soft-agar overlay. The plates were incubated for 16 h at 37°C and the formed plaques were used to enumerate PFU/ml.

### Plasmid construction

All plasmids used in this study are listed in **supplementary table S2**. Plasmids expressing CRISPR guide RNAs were constructed through Golden Gate assembly. First, a GFP dropout plasmid was constructed, expressing GFP upstream of the sgRNA scaffold flanked by BsmBI restriction sites on either side. The desired guide sequences were created by annealing single stranded oligonucleotides, with BsmBI specific overhang^55^. For that, 150 pmol of each oligo was mixed with 0.5 µL T4 polynucleotide kinase (NEB, Cat. # M0201L), 2.5 µL T4 DNA Ligase buffer (NEB, Cat. # M0202L) and adjusted with Milli-Q water to 25 µL total volume. The reaction mix was incubated at 37°C for 30 min, followed by heat inactivation at 65°C for 10 min. Afterwards, the oligos were incubated at 95°C for 5 min, followed by a gradual cooling down by 1°C per 15 sec until it reaches 15°C. For the Golden Gate assembly, 40 fmol of GFP dropout plasmid and 2 pmol of the guide insert were mixed with 0.5 µL T7 DNA-ligase (NEB, Cat. # M0318S), 1 µL T4 ligase buffer (NEB, Cat. # B0202S), 0.5 µL BsmBI-v2 restriction enzyme (NEB, Cat. # R0580L), and Milli-Q water to a final volume of 10 µL. The mix was incubated for 25 cycles of digestion at 37°C for 2 min and ligation at 16°C for 5 min, followed by a final digestion at 37°C for 10 min and heat inactivation at 80°C for 10 min. 5 µL of the reaction mix was used to transform chemically competent *E. coli* TOP10 cells.

To introduce the RT into the guide plasmid, a 3 or 4 fragment Gibson assembly was used, depending on the size of the desired edit. The backbone was created by restriction digestion of the guide plasmid using EcoRV-HF (NEB, Cat. # R3195S). For that, 500 ng of plasmid DNA was mixed with 0.5 µL EcoRV-HF, 2.5 µL 10X rCutSmart Buffer and adjusted with Milli-Q water to 25 µL total volume. The reaction was incubated 1 h at 37°C, after which 1 µL rSAP (NEB, Cat. # M0371S) was added and further incubated 1 h at 37°C. The reaction was then heat inactivated at 65°C for 20 min and cleaned up using the DNA Clean & Concentrator™-5 (Zymo Research, D4014). The homology arms were amplified directly from the phage genome and the desired edits were introduced through the overhangs of the primers, if the edit was smaller than 50 bp. For larger edits, the desired edit was amplified as a separate fragment, with 30 bp overhangs matching the homology arms. The Gibson assembly was performed according to the manufacturer’s instructions (NEB, Cat. # E2621L), using 0.06 pmol of inserts and 0.12 pmol of backbone DNA. 2 µL of the assembled mix was used to transform electrocompetent *E. coli* TOP10 cells.

### *E. coli* competent cell preparation and transformation

Electroporation was used for the transformation of *E. coli*. To prepare electrocompetent cells, individual colonies of *E. coli* were cultured in 3 ml of LB medium and incubated at 37°C, shaking orbitally at 220 rpm, for 16 h. The culture was back diluted to OD_600_ equal to 0.1 in 25 ml of fresh LB medium. The culture was incubated at 37°C and 220 rpm until it reached OD equal to 0.5. The bacteria were centrifuged at 4°C at 5000 xg for 10 min and the supernatant was removed. The pellet was then resuspended in 25 ml ice cold 10% glycerol (VWR, Cat. # J61059.AP) and centrifuged again at 4°C at 5000 xg for 10 min. The supernatant was removed and the pellet was resuspended in 1 ml ice cold 10% glycerol. The suspension was transferred into a 1.5 ml tube and centrifuged at 4°C at 5000 xg for 3 min. The pellet was resuspended in 250 µL cold 10% glycerol and 50 µL aliquots were distributed into sterile eppendorf tubes. For electroporation, 30 ng of each plasmid DNA was added to the electrocompetent bacteria. The mix was transferred to an ice cold 1 mm electroporation cuvette and an 1.8 kV, 250 Ω, 25 µF pulse was applied using the Gene Pulser Xcell Total Electroporation System (BioRad, Cat. # 1652660). Transformed *E. coli* cells were recovered in SOC medium (20 g/L Tryptone, 5 g/L Yeast Extract, 0.5 g/L NaCl, 2.5 mM KCl, 10 mM MgCl2, 20 mM glucose, pH = 7.0) at 37°C for 1 h, shaking orbitally at 220 rpm. 100 μL of the bacterial suspension were plated on LB agar medium containing the appropriate antibiotics and incubated 16 h at 37°C.

### Bacterial genome modification

The *waaD* and *waaG* knockouts were done by Lambda Red recombineering according to the Keio collection, including the primer design and experimental procedure^56–58^. First, MG1655 was transformed with the pKD46 helper plasmid, containing the Lambda Red recombineering system. The transformation was performed by electroporation, as described above. Recovery and incubation was done at 30°C, because the plasmid is heat sensitive. Individual colonies were used to set up cultures in 3 ml LB medium and incubated at 30°C, shaking orbitally at 220 rpm, for 16 h. Afterwards, 100 µL of the culture was backdiluted in 25 ml LB medium containing 0.5% L-arabinose and incubated at 30°C, shaking orbitally at 220 rpm until OD_600_ equal to 0.5 was reached. From there, the bacteria were prepared to be electrocompetent, as described above. 1 µg of linear RT was transformed, the bacteria were recovered in 1 ml SOC medium and incubated at 37°C shaking orbitally at 220 rpm, for 4 h. The recovered bacteria were centrifuged at 5000 xg for 3 min, the supernatant was discarded and the pellet resuspended in 100 µL LB medium. 10 µL of the bacterial suspension was plated on LB plates containing carbenicillin, to confirm the loss of the helper plasmid. The remaining bacteria were plated on LB plates containing kanamycin. Correct integration of the resistance cassette was confirmed through PCR and Sanger sequencing (Microsynth AG). From the confirmed clones, electrocompetent bacteria were prepared as described above, and transformed with the pCP20 helper plasmid encoding an FLP recombinase, using electroporation as described above. Recovery and incubation was done at 30°C, because the plasmid is heat sensitive. Individual colonies were restreaked on LB agar containing carbenicillin, on LB agar containing kanamycin and on LB agar without antibiotics and incubated at 37°C for 16 h. Clones that were sensitive to carbenicillin and kanamycin were screened for the correct excision of the resistance cassette by Sanger sequencing.

### Phage editing assays

For the phage genome editing experiments, we used a two-plasmid setup. The first plasmid contained the nuclease expressed from an arabinose inducible promoter and the second plasmid contained the sgRNA and the RT (homology arms flanking the desired edit). To create editing strains, we co-transformed the two plasmids in *E. coli* MG1655 using electroporation, as described above. We then picked individual colonies to set up 3 ml cultures in LB medium which were incubated at 37°C, shaking orbitally at 220 rpm, for 16 h. Then, the cultures were back diluted to OD_600_ equal to 0.1 in LB medium containing 0.5% L-arabinose (induction medium) to induce the expression of the nuclease. The cultures were incubated at 37°C and 220 rpm until they reached OD_600_ equal to 0.5.

Following, 2 ml bacterial culture aliquots were centrifuged for 3 min at 5000 xg, the supernatant was removed and the pellet was resuspended in 200 µL of induction LB medium. Phage lysate was serially diluted in SM buffer, mixed with 100 µL of condensed bacterial culture and 3 ml of 0.7% soft agar and poured on LB plates. The plates were incubated for 16 h at 37°C. To harvest the phages, 2 ml of SM-buffer was added onto the plates and the soft agar was scraped off and mixed with the SM-buffer, using a sterile pipette tip. The phage-soft agar-buffer mix was transferred to 2 ml tubes and centrifuged for 1 min at 11000 xg to pellet the agar and cell debris. The supernatant containing the phages was transferred to a fresh tube and screened for desired edits. The region containing the edit was PCR amplified by OneTaq® DNA Polymerase (NEB, Cat. # M0482L) using 1 µL of phage lysate as template and the appropriate primers listed in **supplementary table S3**. The amplicon was purified with the NucleoSpin™ Gel and PCR Clean-up Kit (Macherey-Nagel, Cat. # 11992242) and amplicon sequencing was performed by Oxford Nanopore Technology (Plasmidsaurus Inc.).

### Library construction

To create the plasmid to introduce the randomized sequences into the phage genome, the guide plasmid containing the appropriate sgRNA and the corresponding homology arms was used as template for a PCR reaction. The primers contain the randomized sequences in the 5’ overhang, followed by 40 bp that are complementary to the template. The PCR was done eight times, to get enough material. The PCR product was run on an agarose gel and the correct band was excised and purified using the NucleoSpin Gel and PCR Clean-up Kit (Macherey-Nagel, 740609.250S). The purified products were combined and concentrated using the DNA Clean & Concentrator 5 kit (Zymo Research, D4013). 10 µL of concentrated DNA was mixed with 10 µL NEBuilder® HiFi DNA Assembly Master Mix (NEB, E2621L) and incubated 1 h at 50°C. Afterwards, the DNA was cleaned up using ethanol precipitation to get a highly concentrated product^59^. 1 µg of the concentrated DNA was transformed in 8 x 25 µL NEB® 10-beta Electrocompetent *E. coli* (NEB, C3020K), following the manufacturer’s protocol. After 1 h recovery at 37°C and shaking orbitally at 220 rpm, all eight cultures were combined. 50 µL of the combined culture was used to create a 1:10 dilution series, which was spotted on LB plates to quantify the transformation efficiency, yielding a theoretical library coverage of 0.01%. The remaining culture was used to inoculate 150 ml LB medium and incubated 16 h at 37°C and shaking orbitally at 220 rpm. The overnight culture was then used to isolate plasmid DNA using the ZymoPURE II Plasmid Midiprep Kit (Zymo Research, D4201). The library quality was confirmed using Sanger sequencing (Microsynth AG).

For the creation of the editing strain, electrocompetent *E. coli* MG1655 cells carrying the nuclease plasmid, were freshly prepared. 1 µg of library plasmid was used to transform eight replicates by electroporation. After recovery, some of the culture was serially diluted and plated on LB + Carb agar plates, to evaluate the transformation efficiency and to estimate the coverage of the library size. The remaining culture was back diluted in fresh LB medium and incubated 16 h at 37°C and 220 rpm. The next morning, 300 µL of overnight culture was backdiluted in 25 ml fresh LB medium with 0.5% arabinose, to induce nuclease expression. From there, a regular phage editing experiment was performed to introduce the library into the phage genome.

### Phage library construction and enrichment

To introduce the constructed library into the phage genome, 1 µg of the library plasmid was transformed into the editing strain as described above. After recovery, serially diluted transformants were spotted on LB + Carb plates to determine the transformation efficiency. The remaining bacteria were grown in 25 ml fresh LB + Cm + Carb medium for 16 h at 37°C and 220 rpm. After determining the transformation efficiency, 300 µl of the culture was back diluted in 20 ml fresh LB medium with 0.5% arabinose, to induce nuclease expression. The culture was used to perform plaque assays, and the harvested phages were tested for library integration, as described above.

To enrich the phage library for different characteristics, full plate plaque assays were performed on the *waaD* and *waaG* knockout *E. coli* hosts, respectively. Individual plaques were picked and resuspended in 50 µl SM buffer. Full plates were harvested as described above.

### NGS sample preparation

The desired phage genome fragment was first amplified as described above. A second PCR was performed, using KAPA HiFi DNA Polymerase (Roche, KK2601) with 1 µl amplicon DNA as template, 60°C annealing temperature and 18 cycles. The primers of this second DNA were designed to add flanking Illumina sequencing primer sites to the desired phage sequences (**supplementary table S3**). The DNA was purified using NucleoMag NGS Clean-up and Size Select kit (Macherey-Nagel, 744970.5) and used as template for a follow-up PCR to add Illumina unique dual indices (UDI). This PCR was also performed using KAPA HiFi DNA Polymerase with identical parameters but only doing 17 cycles. The DNA was again cleaned up with the NucleoMag NGS Clean-up and Size Select kit.

The samples were pooled and sequenced using a NovaSeqX, with 150 nt paired-end sequencing and 15 million reads per sample.

### Data analysis

The sequencing results were uploaded to the public Galaxy webserver at <u>usegalaxy.eu</u> and a custom workflow was used to determine the editing rates^60–68^. The custom workflows can be accessed using the following links: https://usegalaxy.eu/u/freng/w/freebayes-mutation-calling and https://usegalaxy.eu/u/freng/w/library-analysis-of-ngs-data. The sequence logos were created using the WebLogo 3 tool, by directly inputting the sequence data^69,70^.

Variant efficacy was computed as a log-fold-change between a variant’s output and input read counts. Included variants were present in either of the replicates with at least 5 reads in both input and output sets (1458/1362 for waaDi /waaGi). We modelled efficacy from amino-acid sequences using two models: The additive model assigns each position-residue its mean log-fold-change over the training set and scores a variant as the sum of its per-position effects (or as a mean position effect across all amino acids if not present in the training set). The epistatic model is a support vector regression capturing interactions between positions (from scikit-learn^71^; third-degree polynomial kernel, an optimized regularization parameter C between 0.01 and 1,000, and an optimized kernel coefficient gamma (scale or auto)). Variant efficacies were predicted on held-out variants by a 5-fold cross-validation: train on 80% of variants, predict on the rest, repeat five times to predict for all variants. Model performance was scored using a Spearman correlation, since our goal was to predict the rank order of the variants rather than specific log-fold-change values, which are less biologically relevant in this context. To compare model performance, 95% confidence intervals were obtained by bootstrap resampling of test variants (10,000 resamples, paired across models and replicates^72,73^); model differences were considered significant when the 95% CI of the per-resample difference excluded zero. All analyses are available at https://github.com/JonesLabEU/phagelib.

### RB-TnSeq fitness assay

1 ml aliquot of Keio_ML9 RB-TnSeq^74^ mutant library was thawed and inoculated into 25 ml of LB medium supplemented with kanamycin (50 μg/ml). The culture was incubated at 37°C until it reached an OD_600_ of 0.4–0.6. We then collected cell pellets (1.5 ml x 3) to serve as a common reference for BarSeq, referred to as the time-zero.

The culture was diluted to an OD_600_ of 0.04 in 2 x LB media (350 μl) and then mixed with an equal volume (350 μl) of phages that were diluted in phage dilution buffer (Teknova:S249). This mixture was distributed into a 48-well microplate (700 μl per well) to conduct competitive mutant fitness assays with various T7 phage mutants at a multiplicity of infection (MOI) of 1. We also designated three control “no-phage” competitive mutant fitness assays, in which we substituted the phages with the phage dilution buffer. The microplates were incubated in Agilent BioTek 800 TS absorbance reader with orbital shaking, and OD_600_ readings were taken every 15 minutes for 16 hours. Following the experiment, survivors were collected, pelleted, and genomic DNA was extracted using the QIAamp 96 DNA QIAcube HT Kit (Cat no: 51331), adhering to the manufacturer’s protocol. Following the DNA isolation, we performed the 98°C BarSeq PCR protocol as described previously^74^. The BarSeq PCR products were sequenced on the Illumina HiSeq 4000 instrument using 50 single-end runs. RB-TnSeq fitness data were analysed as previously described^45^.

### Experimental validation of fitness genes by EOP assay

To validate the phage resistance phenotypes identified through the TnSeq assay, we conducted EOP assays on selected *E. coli* mutants from the Keio collection^56^. All mutant strains utilized in this study were confirmed through Sanger sequencing. Phage quantification was performed using the spot titration method. 2 μl of serially diluted phages were spotted onto a solidified lawn comprising approximately 10 ml of 0.5% top agar, which had been inoculated with 200 μl of a fresh overnight bacterial culture, and then incubated overnight at 37°C. The EOP was calculated as the ratio of plaques on the mutant *E. coli* strain to the plaques on the parental WT *E. coli* strain (BW25113). EOPs were derived from a minimum of two biological replicates. The source data file contains all EOP data.

### Data availability

The phage library sequencing data has been deposited in the National Center for Biotechnology Information database and will become available upon acceptance of this manuscript for publication (BioProject accession PRJNA1356711). All data used for this study are provided in the source data file.

## Author contribution

**Methodology**: all authors. **Phage assays**: F.E., S.K.V. **Bioinformatic analysis**: F.E., S.K.V., J.K. **RB**-**TnSeq assays**: S.K.V. **Writing manuscript**: F.E., C.P., S.K.V. **Reviewing and editing manuscript**: all authors. **Figure generation**: F.E., J.K., C.P., S.K.V. **Supervision**: C.P., C.L.B., V.K.M. **Funding acquisition**: V.K.M., C.L.B, C.P..

## Acknowledgments

This work was supported by the European Research Council Consolidator grant 865973 (to C.L.B.). C.P. and S.K.J. acknowledge funding from the Research Council of Lithuania under the Programme University Excellence Initiatives of the Ministry of Education, Science and Sports of the Republic of Lithuania (Measure No. 12-001-01-01-01, Improving the Research and Study Environment), project No: S-A-UEI-23-10. S.K.J. acknowledges funding from the European Research Council, Project 101078247-PROTEGE. V.K.M acknowledges the Biopreparedness Research Virtual Environment (BRaVE) Phage Foundry at Lawrence Berkeley National Laboratory which is based upon work supported by the U.S. Department of Energy, Office of Science, Office of Biological & Environmental Research under contract number DE-AC02-05CH11231.

## Supplementary figures

**Figure S1:**
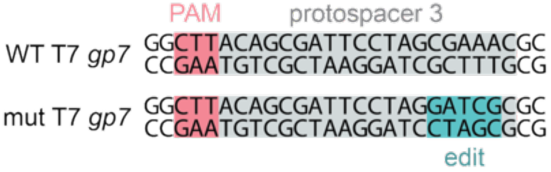
Target site for the T7 *gp7* gene. The target protospacer sequence of the phage T7 *gp7* gene is highlighted in gray, with the ScCas9 PAM highlighted in pink and the introduced edit highlighted in teal.

**Figure S2.**
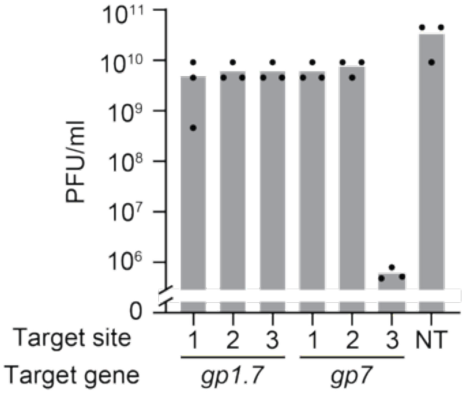
CRISPR-Cas9 counterselection assays for T7 *gp1.7* and *gp7* genes. Phage titers (PFU/ml) resulting from counter-selection by WT ScCas9 targeting three different sites of the genes *gp1.7* and *gp7*, compared to a non-targeting control. Each bar represents the mean of three independent biological replicates.

**Figure S3.**
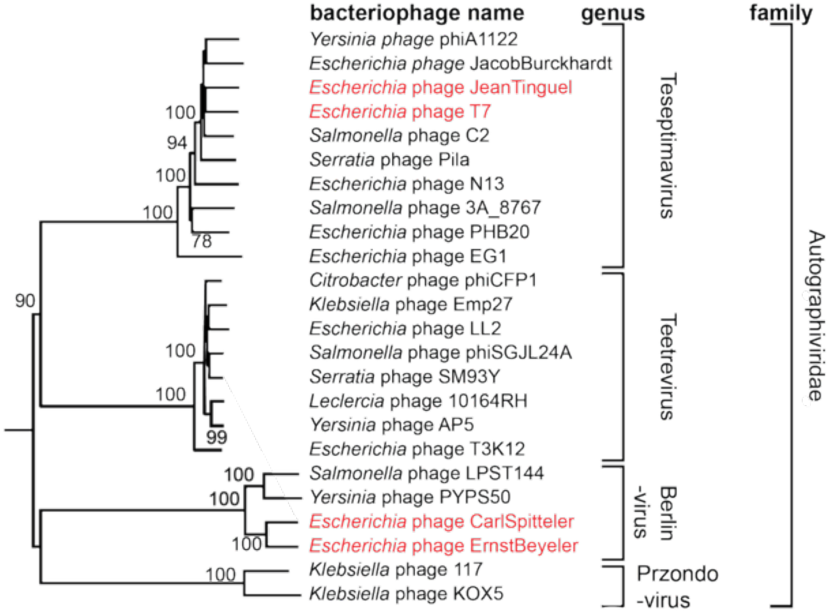
Phylogenetic tree of various related Autographivirdae. Highlighted in red are the phages used in this study (Bas64 = JeanTinguel, Bas67 = CarlSpitteler, Bas68 = ErnstBeyeler). Data adapted from Maffei, E. *et al*., (2021)^75^.

**Figure S4.**
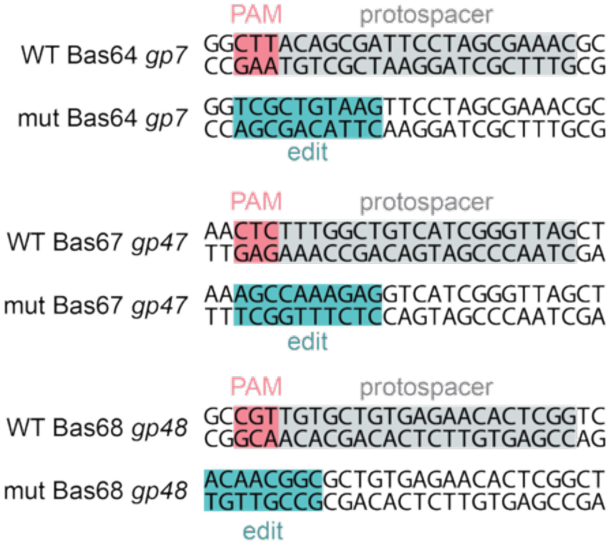
Target sites for Bas64 *gp7*, Bas67 *gp47* and Bas68 *gp48.* Targeted protospacer sequences of phages Bas64 *gp7*, Bas67 *gp47* and Bas68 *gp48* are highlighted in grey, with the ScCas9 PAM highlighted in pink and the introduced edit highlighted in teal.

**Figure S5.**
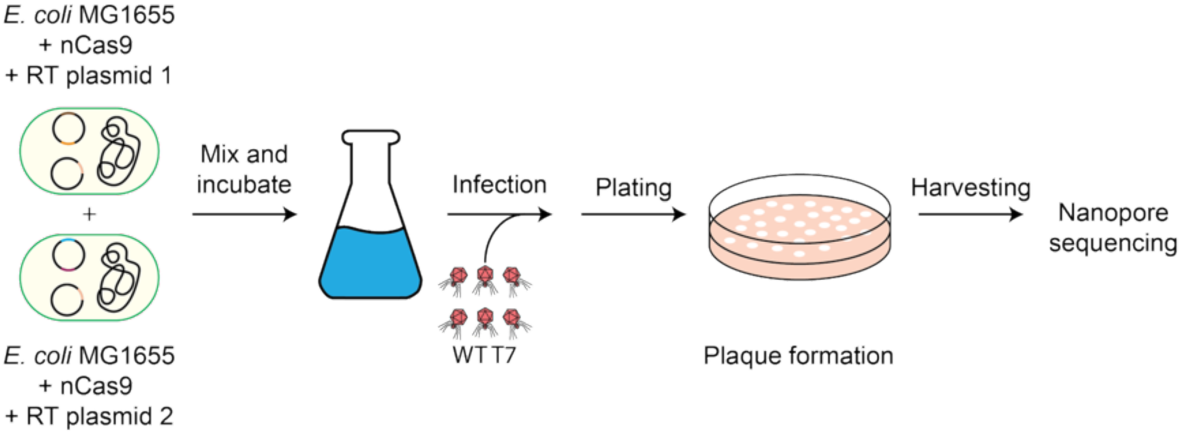
Workflow for multiplexed T7 phage genome editing. Bacterial cultures carrying plasmids encoding for different sgRNAs and RTs are mixed, before infection with WT T7 phages. The resulting plaques containing the different mutant phages were harvested and screened through Oxford Nanopore sequencing.

**Figure S6.**
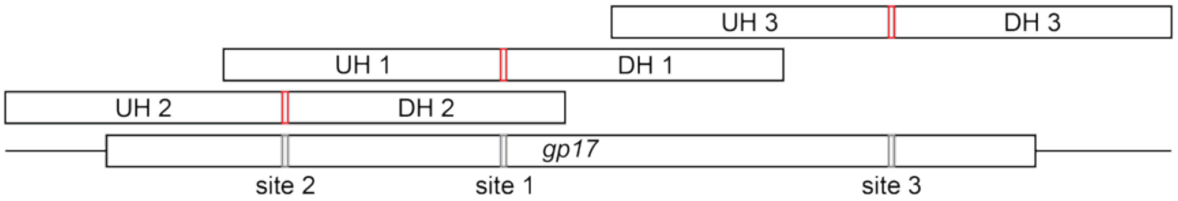
Schematic overview of the three different target sites for multiplexed editing of the T7 *gp17* gene. The area covered by the upstream homology arms (UH) and downstream homology arms (DH) and the target sites are shown. Red boxes indicate the repair sequence flanked by the homology arms.

**Figure S7.**
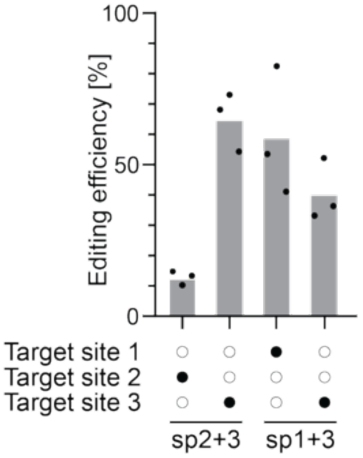
Editing efficiency for the individual target sites after multiplexed editing. Two target sites at the *gp17* gene were targeted simultaneously and editing efficiency of the individual target site is shown. Each bar represents the mean of three independent biological replicates.

**Figure S8.**
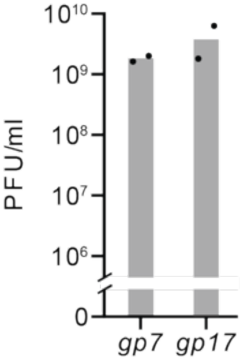
Phage titers resulting from infecting *E. coli* carrying the *gp7* and *gp17* library plasmids with phage T7. Each bar represents the mean of two independent biological replicates.

**Figure S9.**
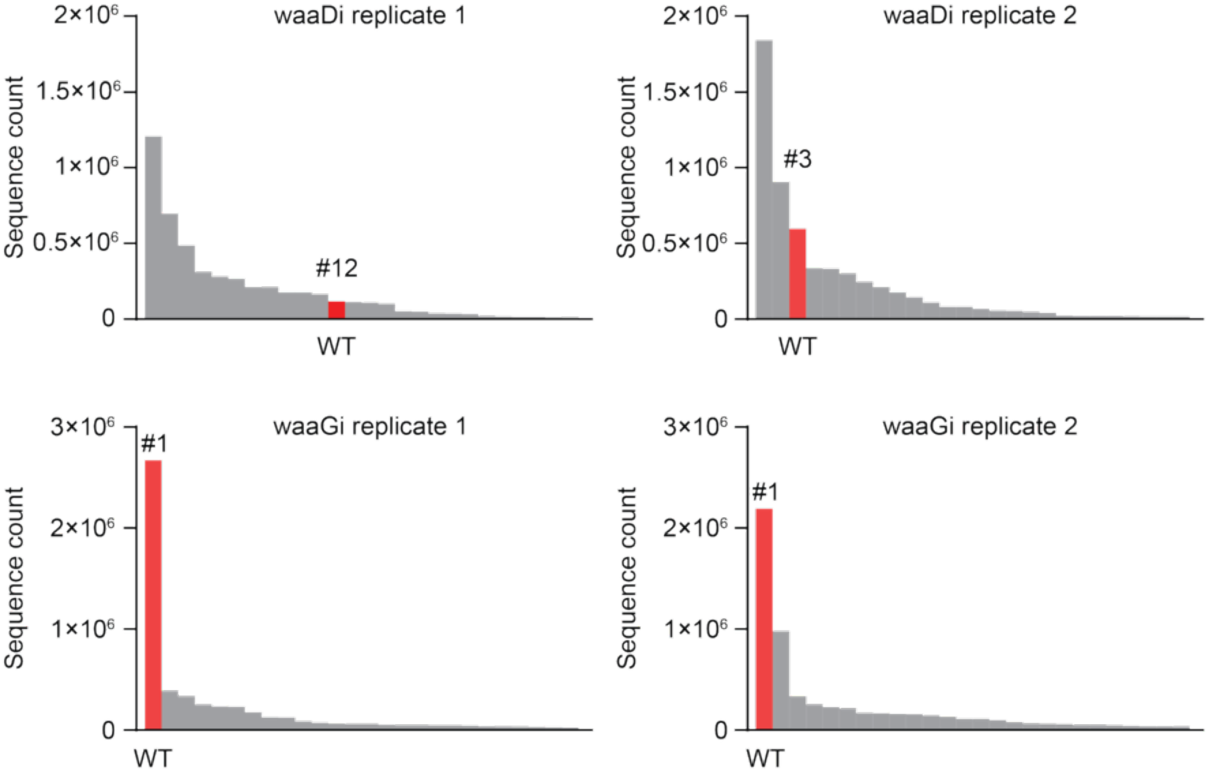
Sequence counts of individual sequences of waaDi and waaGi. The WT sequence is highlighted in red and its ranking based on sequence counts is listed above the bar.

**Figure S10.**
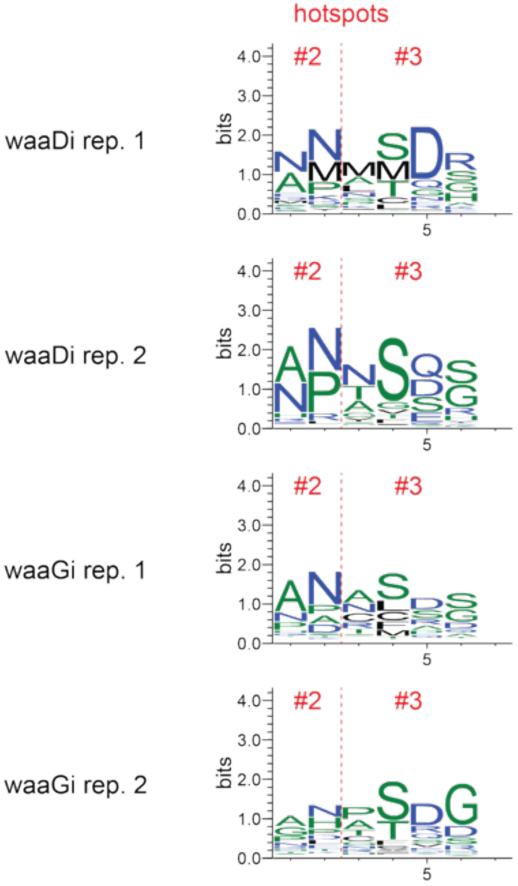
Sequence logos of the waaDi and waaGi mutants. Hotspot #2 (left) and #3 (right) have been separated and WT T7 phage sequences were removed before generating the logos.

**Figure S11.**
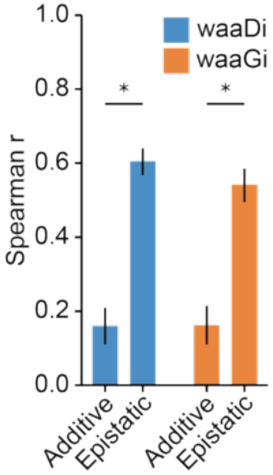
Additive and epistatic model performance for waaDi and waaGi. Bars report a cross-validated correlation between predicted and measured efficacy for the additive and epistatic models cross-validation. Error bars are 95% bootstrap confidence intervals (10,000 resamples). Asterisks: paired bootstrap 95% CI of the model difference excludes 0.

**Figure S12.**
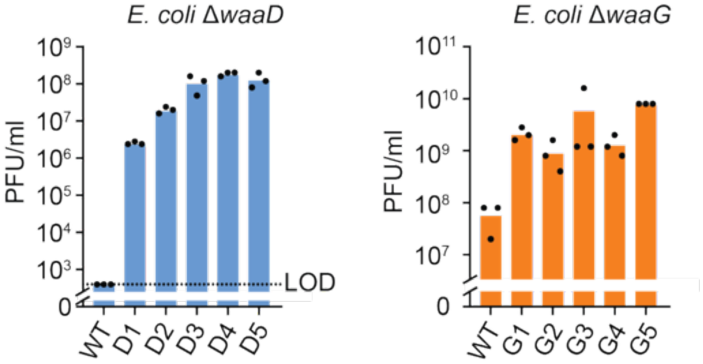
Infectivity of T7 mutants isolated on LPS-deficient *E. coli* hosts. Phage titers of WT T7 and mutant phages isolated from *E. coli* Δ*waaD* (left) and Δ*waaG* (right) determined on the corresponding *E. coli* LPS mutant. Each bar represents the mean of three independent biological replicates.

**Figure S13.**
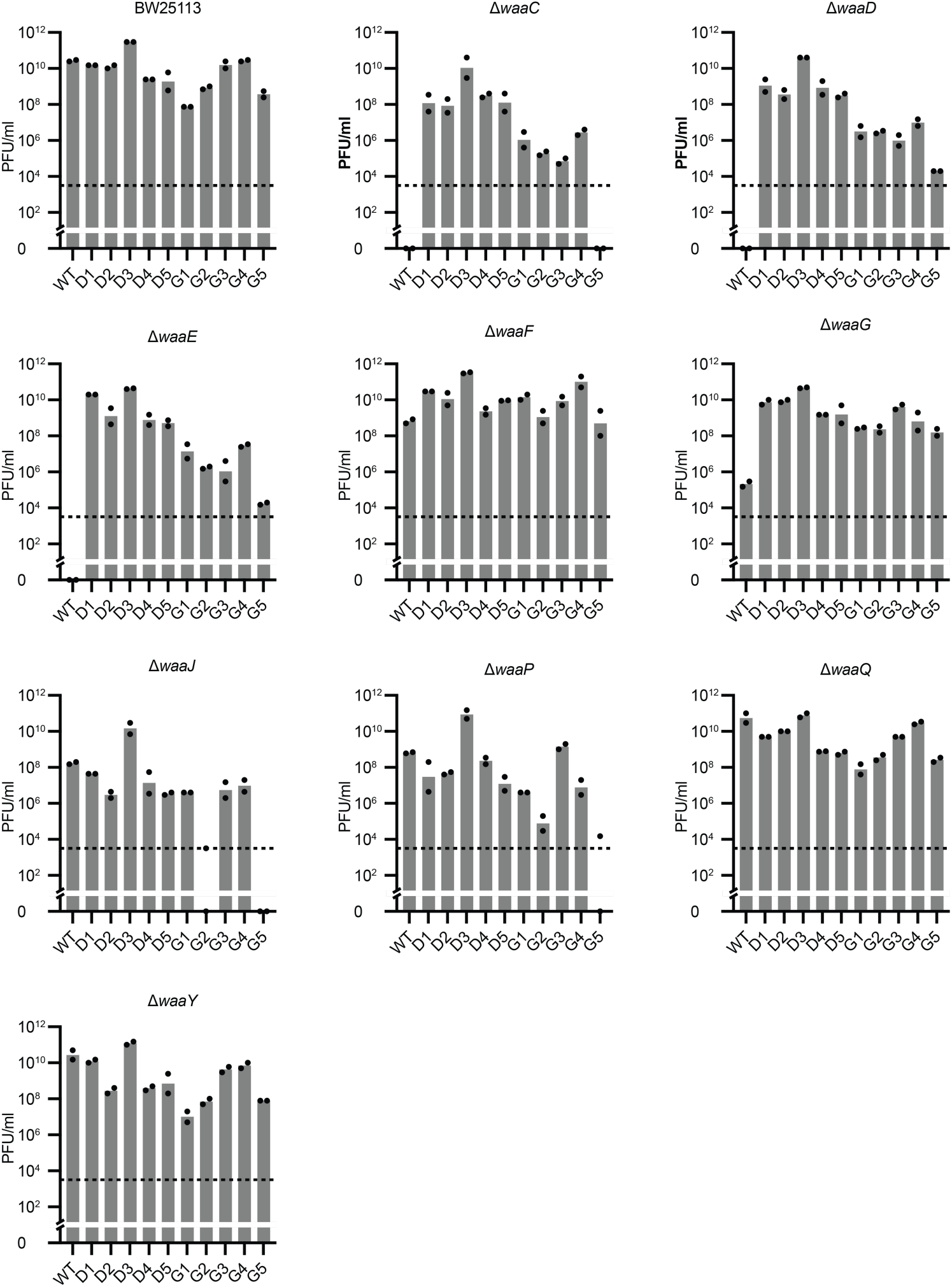
Host-range profiling of T7 mutants across defined *E. coli* LPS mutants – Combined for each mutant host. Phage titers of WT T7 phage and five waaDi and waaGi, determined on WT *E. coli* BW25113 and nine defined *E. coli* BW25113 LPS mutant hosts (Δ*waaC*, Δ*waaD*, Δ*waaE*, Δ*waaF*, Δ*waaG*, Δ*waaJ*, Δ*waaP*, Δ*waaQ*, and Δ*waaY*). Each bar represents a biological duplicate. The dotted line represents the limit of detection.

**Figure S14.**
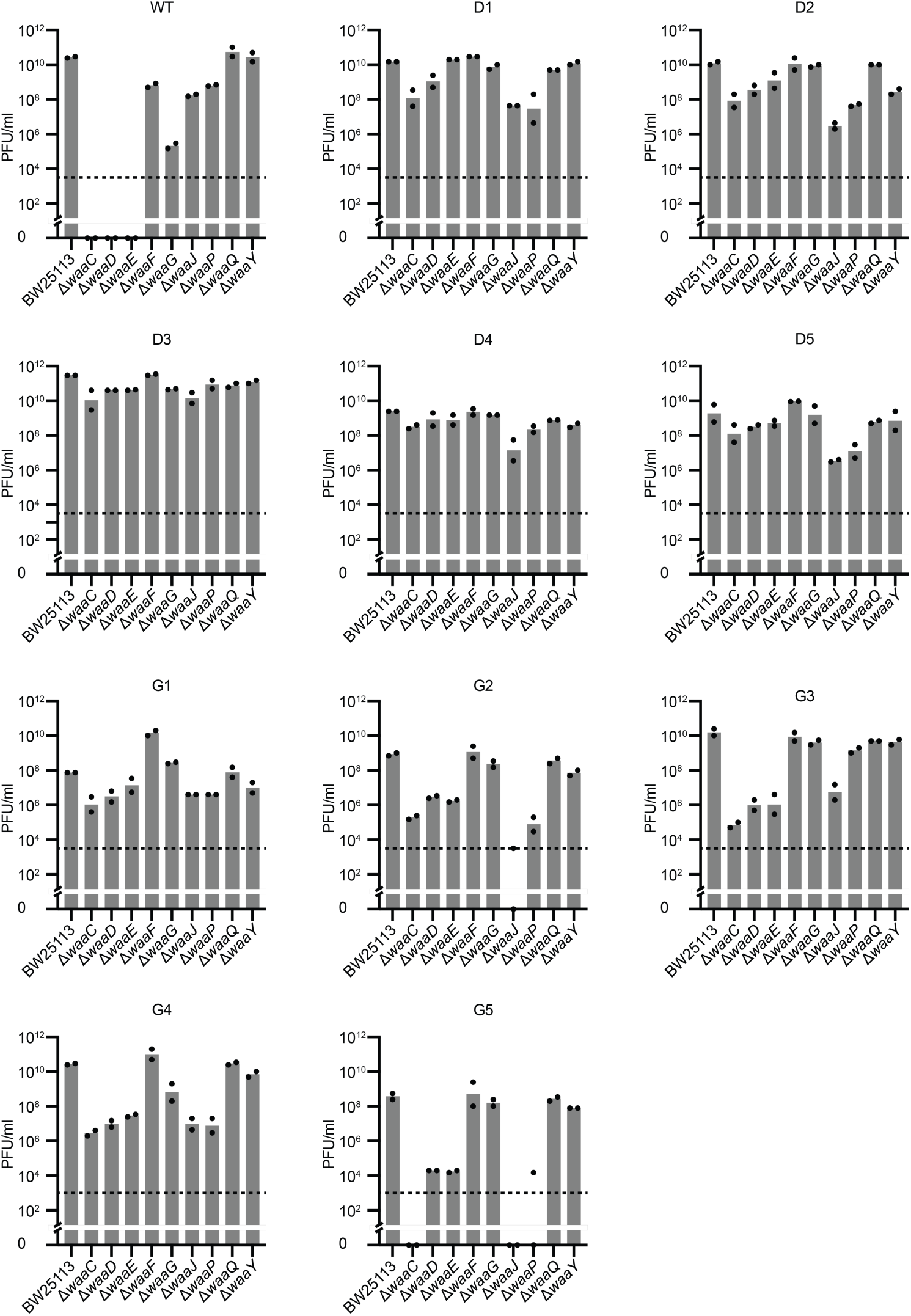
Host-range profiling of T7 mutants across defined *E. coli* LPS mutants – Combined for each T7 isolate. Phage titers of WT T7 phage and five waaDi and waaGi, determined on WT *E. coli* BW25113 and nine defined *E. coli* BW25113 LPS mutant hosts (Δ*waaC*, Δ*waaD*, Δ*waaE*, Δ*waaF*, Δ*waaG*, Δ*waaJ*, Δ*waaP*, Δ*waaQ*, and Δ*waaY*). Each bar represents a biological duplicate. The dotted line represents the limit of detection.

**Figure S15.**
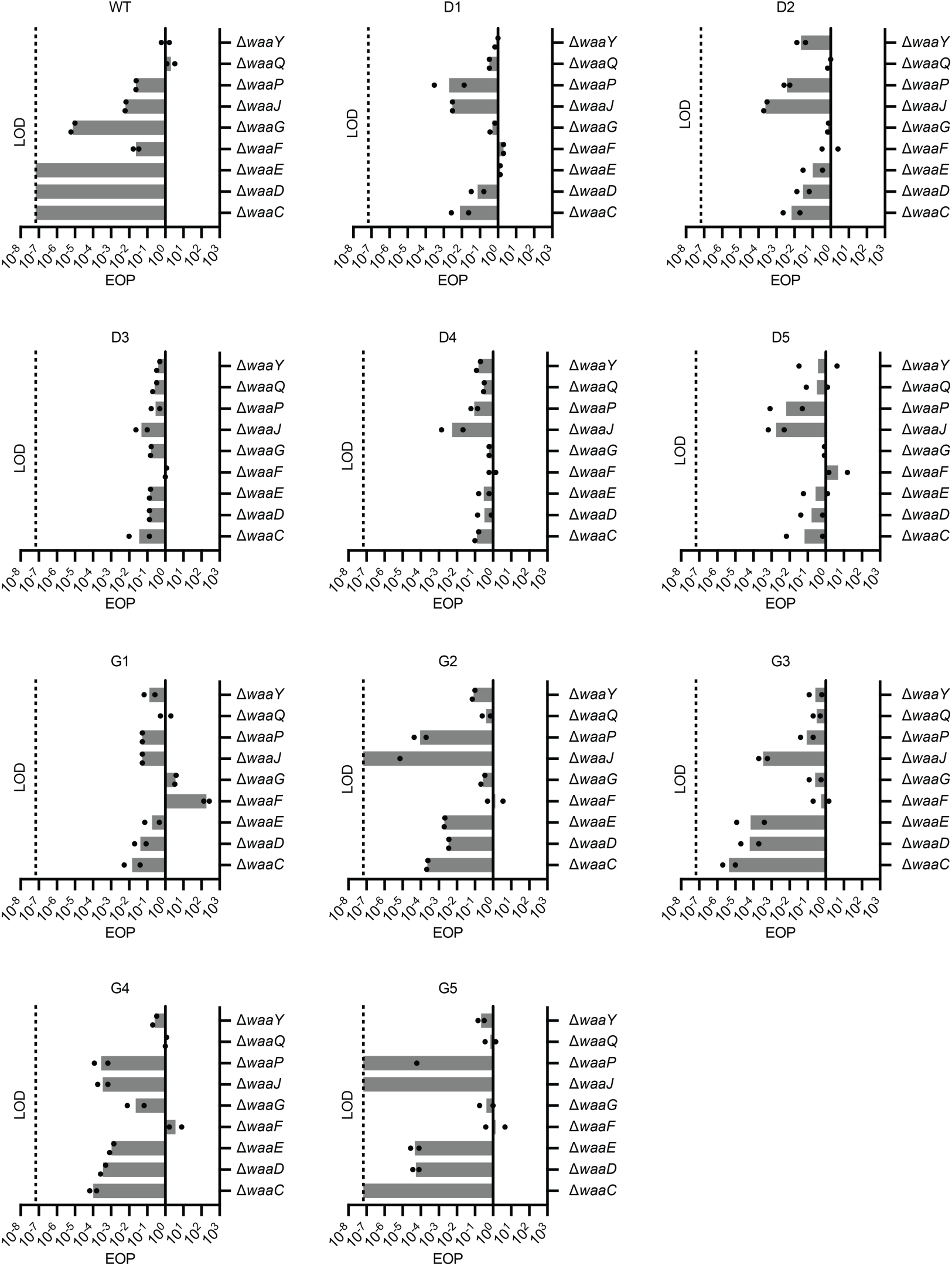
EOP assays against various *E. coli* mutant hosts. EOP of the WT T7 phage and the five isolates selected on Δ*waaD* (D1-D5) and on Δ*waaG* (G1-5) *E. coli* hosts, against nine defined LPS mutant hosts (Δ*waaC*, Δ*waaD*, Δ*waaE*, Δ*waaF*, Δ*waaG*, Δ*waaJ*, Δ*waaP*, Δ*waaQ*, and Δ*waaY*). Data are combined for each T7 phage isolate. All EOP assays were performed in duplicate. LOD: Limit of detection.

